# Learning local geometry and nonlinear topology of neural manifolds via spike-timing dependent plasticity

**DOI:** 10.1101/2025.08.27.672728

**Authors:** Nikolas Schonsheck, Chad Giusti

## Abstract

Neural manifolds are an indispensable framework for understanding information encoded by activity in neural populations. While some neural manifolds are linear and can be recovered from population activity using standard techniques, many neural manifolds exhibit nonlinear global topology for which such tools can be less effective. Notably, circular and toroidal manifolds describe activity in neural systems across a range of species; common examples include orientation-selective simple cells in primary visual cortex, head-direction cells in thalamic circuits, and grid cells in entorhinal cortex. That such structured information appears in both sensory and deep-brain regions raises a basic question: is the propagation of nonlinear coordinate systems a generic feature of biological neural networks, or must this be learned? If learning is necessary, how does it occur? In this paper, we apply methods from topological data analysis developed to quantitatively measure propagation of such nonlinear manifolds across populations to address these problems. We identify a collection of connectivity and parameter regimes for feed-forward networks in which learning is required, and demonstrate that simple Hebbian spike-timing dependent plasticity reorganizes such networks to correctly propagate circular coordinate systems. We observe during this learning process the emergence of geometrically non-local experimentally observed receptive field types: bimodally-tuned head-direction cells and cells with spatially periodic, band-like receptive fields. These observations provide quantitative support for the hypothesis that simple biologically plausible plasticity mechanisms suffice to induce changes in the structure of neural architectures sufficient to explain the appearance of such features in real neural systems.

## Introduction

Experimental observations of *in vivo* neural populations demonstrate that nonlinear neural manifolds arise in multiple brain regions across many species. The most broadly reported of these are circular coordinate systems and tori, which represent multiple independent circular coordinates. Important examples include head direction (HD) cells (reported in mammals [1, 2, 3], birds [4], and insects [5]), simple cells (reported in primary visual cortex of cats [6, 7] and nonhuman primates [8]), and grid cells (reported in entorhinal cortex of many mammalian species [9, 10, 11, 12]). Notably, each of these systems participates in a hierarchy of neural information integration: simple cells in primary visual cortex project to higher visual cortical areas [13, 14, 15, 16] and it is known that parahippocampal circuits containing HD cells and entorhinal cortex with grid cells participate in a number of feedback loops [17, 18, 19, 20, 21, 22, 23, 24]. This suggests that such nonlinear organization is not simply a local feature of neural populations, but also must be propagated to, and thus learned by, downstream brain regions.

To study how circular coordinate systems arise in and propagate through neural circuits, we must be able to reliably detect and quantify circular topology in population-level neural activity. Many nonlinear dimensionality reduction techniques have been developed [25, 26, 27] to understand the latent structure of high-dimensional data. However, such methods generally are non-deterministic, sensitive to parameter choice, and produce embeddings that are difficult to test for faithfulness to high-dimensional structure. As illustrated in Fig 1(e) they can fail to detect true nonlinear structure in population activity, or falsely suggest the presence of such structure. As these methods do not carry theoretical guarantees or reliable quantitative measures of goodness of fit to global topology, these failures can go undetected in settings where the ground truth is unknown. On the other hand, tuning curves—which are experimentally accessible in the context of known stimuli—tell us only about the properties of single units and not about how populations of neurons jointly encode structure. Analyzing tuning curves alone can thus similarly lead to incorrect conclusions.

**Figure 1.**
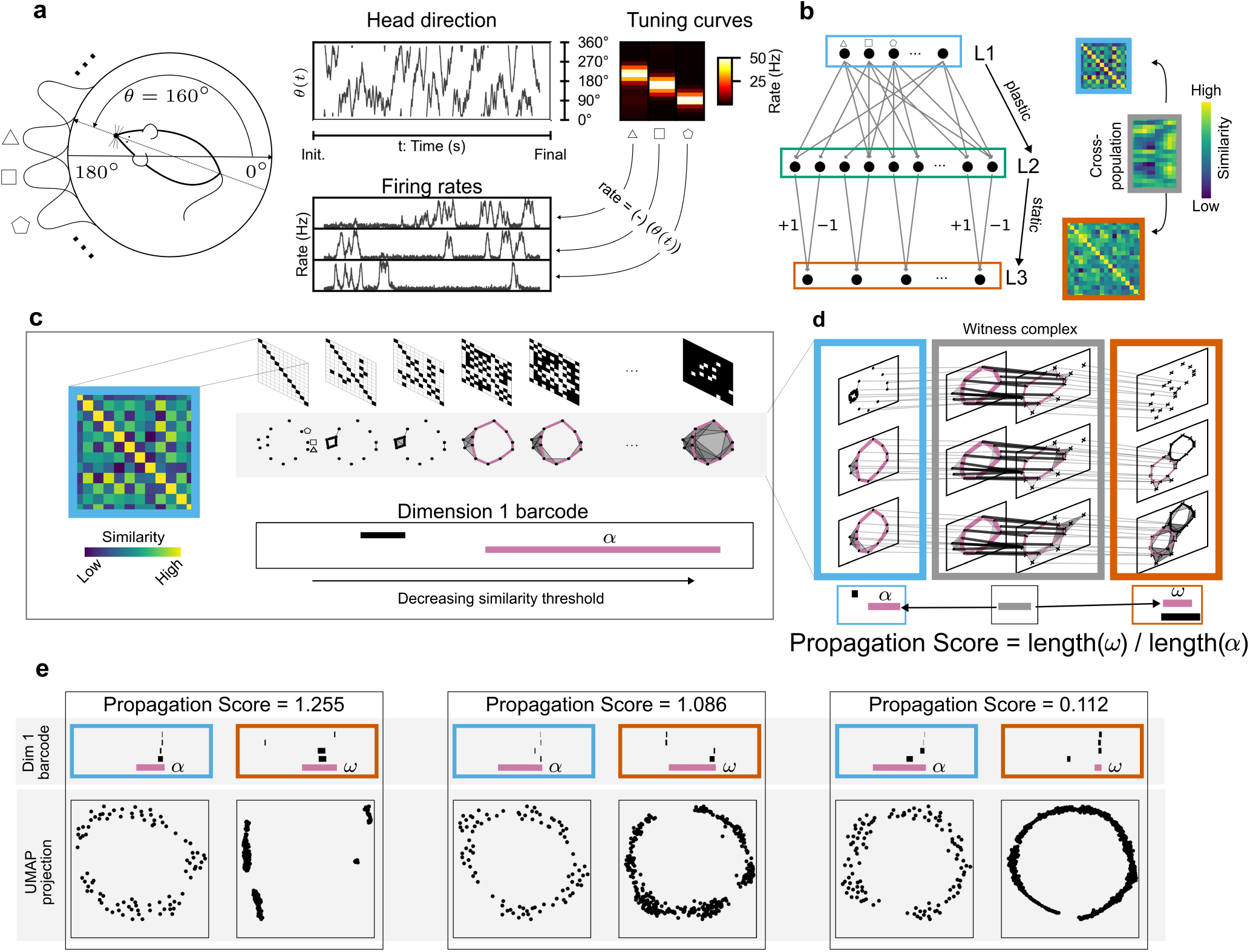
Tracking circular coordinate systems through a multilayer feedforward network using barcodes. **(a)**, Firing rate rasters of a population of synthetic head direction cells are constructed using a simulated heading stimulus and tuning curves extracted from extracellular recordings in freely moving mice (see *Methods and Materials*). **(b)**, (Left) Architecture of the model network: L1 comprises Dale neurons with excitatory projections to L2; projections from L2 to L3 are constrained so that each L3 neuron receives one excitatory and one inhibitory signal. (Right) For a fixed set of synaptic weights and heading stimuli, intrapopulation similarity matrices are computed for populations L1 (blue) and L3 (orange) using cross-correlograms of the associated firing rate rasters; cross-population similarity between L1 and L3 is similarly computed (grey). **(c)**, (Top row) Given a symmetric similarity matrix (blue outline), a sequence of binary adjacency matrices is constructed by systematically decreasing a similarity threshold. (Middle row) Each binary adjacency matrix determines a *clique complex*, a generalization of a graph that natively encodes multi-unit similarity. (Bottom row) A summary statistic that tracks the evolution of topological structure called a *barcode* is computed from the sequence of clique complexes. Long bars (pink) indicate robust circular topology in the L1 population encoding. **(d)** The *method of analogous cycles (MAC)* [32] is used to determine if topological structure detected downstream (orange outline) encodes the same topological information as *α*. If a match is found (pink), the robustness of propagation is quantified by comparing lengths of matched bars (*α* and *ω*, respectively). **(e)**, Standard dimensionality reduction techniques such as UMAP are unreliable for detecting matched circular coordinates across populations. (Top) L1 and L3 barcodes for a sample network with matches reported by MAC highlighted in pink. (Bottom) UMAP projections of the correlation matrix. (Left) a false negative, (Middle) a true positive, (Right) a false positive.

To overcome these limitations, we observe that despite the varied details of the individual neural systems discussed above, they can be represented using a common mathematical framework: the stimulus space model [28]. In this model, neurons and stimuli correspond to points in an underlying metric space in which neurons have receptive fields, understood as geometrically localized regions in the space; populations with circular coordinate systems trace out circular regions in the stimulus space. This framework dovetails with computational tools from topological data analysis [29, 30, 31] which provide techniques for detecting and quantifying circular topologies, including a newly developed technique called the *method of analogous cycles* (MAC) [32] for matching topological structure across neural manifolds without reference to external stimuli or behavioral correlates. Combining these techniques, we obtain a mathematically well-founded and quantitatively falsifiable framework for investigating mechanisms through which neural populations can learn to represent circular coordinate systems.

A number of algorithms for learning geometric structure have been described in the literature [33, 34, 35, 36]. However, these are largely formulated in the context of problems in machine learning or high-dimensional data analysis, and therefore rely on biologically implausible mechanisms such as gradient backpropagation. Our goal is to understand the emergence of such structure in biological neural networks, which are invariably noisy and may include high levels of inhibition. Therefore, we are led to ask: is there a simple and biophysically grounded mechanism that is sufficient for the learning of such nonlinear neural manifolds? Two prior results motivate our approach. First, spike-timing dependent plasticity (STDP) [37, 38, 39] can act as a competitive Hebbian learning mechanism [40]. Second, a classical result in manifold learning [41] suggests that Hebbian learning can induce the formation of global topology by assembling local geometry. We therefore hypothesized that classical Hebbian STDP can induce learning of nonlinear neural manifolds, including circular and toroidal coordinate systems.

In this paper, we investigate this hypothesis using simulated feed-forward networks equipped with Hebbian STDP. We model biologically plausible neuronal responses for neurons in the input layer using tuning curves and rate maps extracted from extracellular recordings of head-direction cells in mice [2, 42] and grid cells in rats [43]. We then introduce a new quantiative measure of effective propagation of topological structure. This statistic allows us to identify a range of biologically realistic architectures and parameter regimes in which learning is required to propagate circular coordinate systems, and validate that Hebbian STDP suffices for such networks to learn global circular and toroidal neural manifolds. Finally, we investigate how STDP-induced learning shapes downstream tuning profiles through the interaction of excitatory and inhibitory connectivity. In this context, we observe the spontaneous emergence of experimentally observed phenomena such as bimodal head direction cells and cells with receptive fields in the form of spatially periodic bands, first reported in [44]. We leverage this analysis to provide potential mechanistic explanations for the presence of these phenomena in biological neural networks.

## Results

To investigate the propagation of circular coordinate systems, we utilized a feedforward network architecture which minimizes selections of parameters subject to the following necessary constraints. First, the architecture needs to allow circular coordinate systems to be enforced in the input layer. Second, the network must be capable of modeling standard biological features such as sparsity and excitatory/inhibitory (E/I) balance. Third, it must admit STDP as a method of updating weights for both excitatory and inhibitory neurons. A robust but simple model which satisfies these assumptions can be constructed using three layers of standard ReLU rate-model neurons. Neurons in the first layer (L1) are excitatory Dale neurons. These project via an arbitrary weight matrix *W* to a second layer (L2) consisting of equally-sized populations of inhibitory and excitatory rate-model neurons. Neurons in the second layer are partitioned into pairs of excitatory and inhibitory neurons, and each pair projects to a distinct readout neuron in the third layer (L3) with fixed unit weights (Fig 1(b)). Associating tuning curves that model response to specific stimuli (e.g., head direction or location) to individual L1 neurons induces the corresponding topology in the L1 population activity. Selecting an appropriate matrix *W* models network features such as sparsity and E/I balance. Finally, sampling from an inhomogeneous Poisson process using the computed firing rates of individual neurons simulates the spiking with which we apply Hebbian STDP to update *W*.

For a fixed weight matrix *W*, we measured the network’s ability to propagate circular coordinate systems by modeling L1 response to a simulated stimulus and then passing this activity through the network (Fig 1 (a) and (b)). Then, we computed pairwise dissimilarity matrices for the activity of L1 and L3 populations, and for L1 to L3 interpopulation dissimilarity (Fig 1 (b) blue, orange, and grey respectively). We used persistent homology to quantify the presence or absence of circular topology in the population activity (Fig 1(c)). Lastly, we employed MAC [32] to determine whether the enforced circular structure in L1 matched to any structure detected in L3, and quantified the robustness of propagation by computing the ratio of the length of the bars corresponding to the matched features (Fig 1(d)). We refer to this procedure as a *standard propagation trial* (see *Materials and Methods* for further details).

### Learning a single circular coordinate system with STDP

In order to investigate how nonlinear coordinate systems propagate and form, we began with the simplest such structure, a single circular coordinate system. To enforce this structure in L1 population activity, we modeled individual neural response as a function of heading direction using tuning curves extracted from extracellular recordings of head-direction cells in mice. A three-layer network (Figure 1) was randomly initialized and a standard propagation trial was conducted to determine whether the circular topology would propagate downstream to L3 through a randomly-wired network. The network architecture was determined by 7 different hyperparameters, including excitatory/inhibitory balance in the L1→ L2 layer and noise in the L1 encoding. We tested 648 distinct hyperparameters combinations, each over 10 trials (see *Materials and Methods*).

For 418 of the 648 (64.51%) architectures tested, the mean Propagation Score (PS) over 10 trials was below 0.25, indicating in these cases that circular structure was not reliably preserved downstream. Propagation failure was particularly pronounced in the presence of net inhibition: among the 216 architectures with inhibitory bias in the L1 to L2 initialization, 181 (83.80%) yielded mean PS below 0.25. In the most extreme cases, with net inhibition and nonzero levels of noise introduced to the L1 encoding by including neurons with no obvious circular tuning (see *Materials and Methods*), 137 of 144 hyperparameter combinations (95.14%) had mean PS below 0.25. These results suggest that randomly initialized networks, particularly those with net inhibition and noise in the initial encoding, will not reliably propagate a circular coordinate system. Given the inherent noise in neural coding and that inhibition dominates in many neural circuits [45, 46, 47, 48, 24], this raises a question: can such networks to learn to propagate circular coordinate systems via known biophysical mechanisms?

To answer this, we began with the same set of 6480 networks and simulated 5000 seconds (~83 minutes) of open-field exploration during which we updated synaptic weights between L1 and L2 neurons over 1-second epochs according to the spike-timing dependent plasticity (STDP) equations in [39, Fig 1.] (see Figure 2 (a)-(b)). For each L2 neuron (and 1s epoch), we scaled the calculated STDP-induced weight changes in two ways. First, in order to prevent tonically high-firing neurons from dominating learning, we amplified plasticity of synapses involving presynaptic neurons that showed high variability in firing rate compared to the previous 10s (see *Methods and Materials* for further information). Second, to maintain network homeostasis, we enforced a maximum synaptic weight of +1 and uniformly scaled all input weights to each L2 neuron to preserve total synaptic weight projecting to the neuron throughout learning; this is in line with known mechanisms of homeostatic synaptic plasticity [49, 50, 51, 52, 53, 54]. After this learning period, we assessed how the changes in network architecture from STDP affected propagation of the circular coordinate by performing a propagation trial with the same stimulus used before network updates.

**Figure 2.**
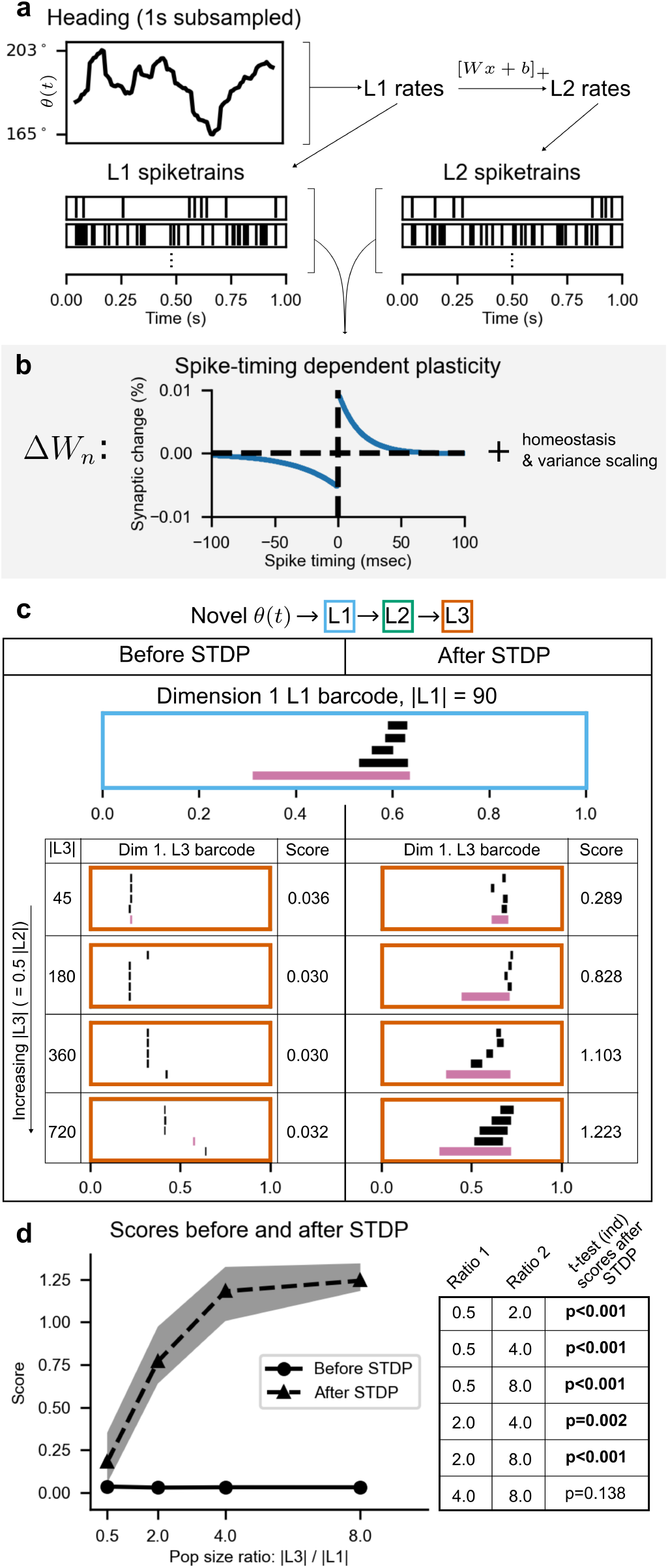
Hebbian spike-timing dependent plasticity supports propagation of a circular coordinate system. **(a)**, Firing rates for L1 and L2 neurons simulated in 1 second learning epochs in response to heading stimuli are converted to spiketrains using an inhomogeneous Poisson process **(b)**, For each epoch, synaptic weights between each pre- and post-synaptic neuron are updated by applying and scaling the Hebbian STDP equations found in [39, Fig 1.] using the respective spiketrains (see *Methods and Materials*). **(c)**, After a learning period of 10^5^ epochs (~1.38 hours simulation time), a novel heading stimulus modeling 25 minutes of freely moving behavior is presented to the network. Downstream encoding of the circular coordinate system before and after STDP learning is assessed using MAC (see Fig. 1, *Methods and Materials*). Robustness of propagation increases with L3 population size. **(d)** Summary of PS before and after STDP learning over 10 trials for each L1 and L3 population size pairing.

We performed paired one-tailed t-tests on PS scores before and after STDP updates on each hyperparameter selection (10 trials each before and after) and found that 511 of the 648 (78.9%) combinations showed a significant (*p* < 0.05) increase in PS after learning. This increase was especially pronounced in the case of inhibition and noise: 132 of 216 (61.1%) architectures with net inhibition and 97 of 144 (67.4%) architectures with both net inhibition and noise yielded PS above 0.75 after learning. These results suggest that simple STDP-modulated updates to synaptic weights can be sufficient to induce downstream learning of a circular coordinate system. However, the variability of the outcomes led us to ask: are there particular features of the network which determine how robustly the target circular coordinate propagates after learning?

### Relative population size modulates learnability of a circular coordinate system

Investigating the results of our experiments, we qualitatively observed that larger downstream population sizes (L2 and L3) were often associated with improved post-learning PS (Fig 2 (c-d)). To quantify this observation, we fit a decision tree model to predict post-learning PS based on the seven model hyperparameters and computed permutation-based importance scores of each hyperparameter. The model provided a reasonable fit to the data (*R*^2^ = 0.35) and revealed three key contributors: the population size ratio |L3| */* |L1| (Δ*R*^2^ = 0.50); excitatory/inhibitory balance (Δ*R*^2^ = 0.21); and connection density (Δ*R*^2^ = 0.07). The measured Δ*R*^2^ for all other hyperparameters was 0.0. This suggests that relative population size is a significant factor in determining propagation success after STDP-induced learning.

To further investigate this effect, we computed Spearman’s rank correlation across all trials to determine how reliably post-learning PS increased with the population size ratio |L3| */* |L1| (see Fig 2 (d)). The effect was most pronounced in the case of inhibitory bias: 40 of 54 configurations with inhibitory bias showed a significant (*p* < 0.05) positive correlation between population size ratio and post-learning PS. Fig 2(d) shows one such configuration. The effect persisted, though diminished, in the case of excitatory/inhibitory balance and excitatory bias: 33 of 54 and 18 of 54 configurations, respectively, showed significant positive correlation. This suggests that learning global topology via STDP is facilitated by a sufficiently high-dimensional ambient neural activity space. Having confirmed that STDP can induce such learning and identified key factors in its success, we wished to understand why this relative population size was important. Based on the results of [41], we expected that that downstream neurons would learn to represent local neighborhoods of the upstream circular stimulus space. We therefore hypothesized that this relative population size phenomenon occurs because a high-dimensional space can accommodate simultaneous learning of sufficiently many independent local receptive fields to cover the space. To determine whether this was the case, we next asked *how* synaptic weights reorganize via STDP-modulated learning to allow propagation of a circular coordinate.

### An emergent process of learning local geometry

In order to propagate local geometry through the network, we expected that the synaptic weights projecting to a given L2, initially randomly distributed across L1 neurons, would concentrate to a subset of L1 neurons in a small arc of the circular stimulus space, while weights to other neurons would decrease. To test this hypothesis, we assigned each L1 neuron a location in a fixed circular stimulus space given by the angle at which its tuning curve peaked. Given a fixed network, we defined, for each L2 neuron *i*, a random variable *X*_*i*_ taking values in the circle, hereafter denoted *S*^1^, determined by the weights and corresponding *S*^1^-valued L1 neuron locations of all inputs projecting to *i*. We then drew an empirical sample from *X*_*i*_ and estimated its probability density function using kernel density estimation with a von Mises kernel (see *Materials and Methods* for further detail). This yielded a density function *f*_*i*_: *S*^1^ *→* ℝ with 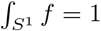, representing the relative distribution of synaptic weight across *S*^1^ projecting to neuron Fig 2(a) shows *f*_*i*_ for a representative L2 neuron before and after network updates. Initially, as illustrated in the upper plot, synaptic weight was distributed more or less uniformly across *S*^1^. As the network reorganized, synaptic weight into this L2 neuron concentrated in a neighborhood of *S*^1^ centered at approximately 100° as shown in the lower plot. This led to the natural question: what effect does this reorganization have on the ability of the network to propagate a circular coordinate?

To answer this question, we fixed a representative network architecture (|L1| = 120, |L3| = 240, moderate noise, inhibitory bias, 50% connection density) and evaluated two metrics across the 10 previously-conducted trails: 1) the average maximum of *f*_*i*_, denoted ⟨max *f*_*i*_⟩ for each L2 neuron *i*, and 2) the PS of the network, both at intervals of 100s over the 5000-second learning period. We observed a reasonably strong correlation (*R*^2^ = 0.536) between ⟨max *f*_*i*_⟩ and PS throughout learning (Fig. 2b). Is there a causal relationship between the two?

To test this, we performed a geometric ablation study on each network after the full learning period by randomly shuffling the synaptic weights projecting to each L2 neuron (10 shuffle trials per network) and recomputing ⟨max *f*_*i*_⟩ and PS. As expected, ⟨max *f*_*i*_⟩ decreased (Fig 2(c), left, student’s *t*-test *p* < 1×10^−4^) and—consistent with our hypothesis—PS also significant decreased (student’s *t*-test *p* < 1×10^−5^). Interestingly, the shuffled PS remained higher than PS at epoch 0, suggesting that non-geometric changes in synaptic strength (neuron-specific bias) also contribute to learning.

Up until this point, our analyses had not differentiated between the excitatory and inhibitory L2 subpopulations. Hence, signals from both become more localized via STDP. However, the functions of the two populations in the network are very different, which led us to ask: what is the relative contribution of each population to the learning process?

To investigate this question, we re-ran the STDP protocol on all network architectures for a fixed choice of L1 and L3 population size (90 and 360, respectively) under three conditions: 1) synaptic weights and threshold for all L2 neurons were updated (control), 2) only inhibitory neurons were updated, and 3) only excitatory neurons were updated. We found that updates to both populations significantly impacted learning, with the excitatory population playing a somewhat more important role. Over 54 network architectures (10 trials each), the mean PS after learning was significantly higher (paired t-test *p* < 0.05) in 53 architectures when both populations were updated compared to when only the inhibitory population was updated, and in 23 architectures compared to when only the excitatory population was updated. Interestingly, in 4 of the 54 architectures, the mean PS was higher after updating only the excitatory population. Together, these results indicate that localization of both the excitatory and inhibitory signal play critical roles in shaping downstream learning.

Our results show how STDP induces a concentration of synaptic weight in downstream neurons to local neighborhoods of the upstream circular stimulus space, and that this reorganization is a key component of a network’s ability to propagate a circular coordinate system. In practice, however, synaptic weights are difficult to directly measure. Can evidence of this reorganization be found in more experimentally-observable properties of downstream neurons?

### Tuning properties of downstream neurons

To investigate how learning reshapes tuning properties, we computed rate curves for L2 and L3 neurons. For L2 neurons, we found a close correspondence between *f*_*i*_ and the corresponding rate curve (e.g., Fig 3(e)), supporting the idea that the geometry of synaptic weight distribution shapes downstream tuning.

**Figure 3.**
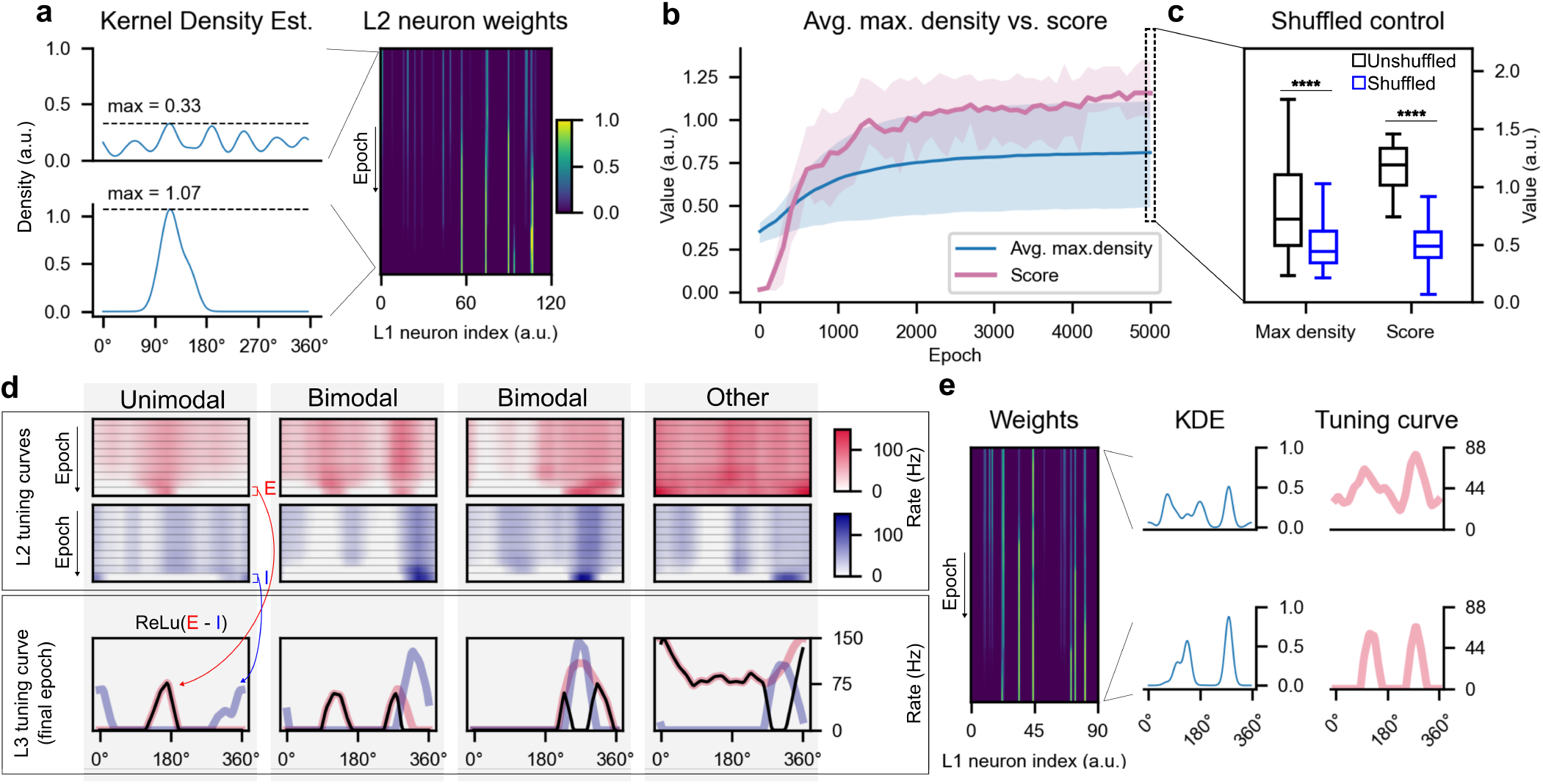
Circular coordinate systems form downstream through an emergent process of learning local geometry. **(a)**, (Right) Synaptic weights from L1 population into a fixed L2 neuron throughout learning. (Left) Kernel density estimates of a synaptic-weight-defined random variable on the circular stimulus space, shown at initial and final learning epochs (see *Materials and Methods*). A higher peak after learning indicates a concentration of synaptic weight in a localized region of the circular stimulus space encoded by the L1 population. **(b)**, PS and average maximum density of L2 neurons are positively correlated over the course of learning and stabilize after approximately 2000 learning epochs. **(c)**, A geometric ablation study is performed by shuffling synaptic weights of L2 neurons and average maximum density and PS is recomputed. Shuffling significantly reduces average maximum density and PS (Welch’s t-test *p* < 1×10^−5^ for both comparisons) demonstrating that geometric organization is necessary for propagation. **(d)**, Evolution of tuning curves of representative L2 and L3 neurons during learning. (Left) A unimodal L3 tuning curve emerges from offset unimodal L2 excitatory and inhibitory curves. (Left middle) A bimodal L3 tuning curve emerges from a bimodal excitatory L2 curve. A bimodal L3 tuning curve emerges from overlapping unimodal L2 excitatory and inhibitory curves. (Right) A non-spatially-tuned L3 neuron emerges. **(e)**, (Left) Weights into excitatory L2 neuron of middle left column of (d). (Middle and right) associated *f*_*i*_ and tuning curves of the neuron before and after STDP.

As expected, many L2 and L3 neurons developed unimodal tuning, while some failed to develop any clear tuning (Fig 3(d), left and right columns, respectively). Surprisingly, we also observed a number of bimodally-tuned downstream neurons. The middle two columns of Fig 3(d) show two such examples. In one case (third column) the bimodal tuning of the L3 neuron is a simple coincidence of the relative alignment of two unimodal curves. In the other case (second column), however, the bimodal tuning of the L3 neuron is inherited from the bimodal tuning of an upstream L2 neuron. To determine whether this bimodal response simply reflected input from bimodal neurons upstream, we computed *f*_*i*_ for this L2 neuron and found two distinct peaks (Fig 3(e)). This indicates that the learning of local geometry can lead to the *emergence* of bimodal tuning, representing multiple neighborhoods of a circular stimulus space, even when no bimodal responses were present in the initial encoding.

Our results thus far suggest that learning of local geometry may be a generic feature of STDP in the presence of a single circular coordinate system in the initial encoding. Does the same phenomenon, in fact, emerge when the input has more complex topology?

### Learning multiple circular coordinate systems with Hebbian STDP

To determine whether STDP can induce propagation of multiple independent circular coordinate systems, we next investigated the same STDP learning process using L1 neurons equipped with 2-dimensional rate maps imputed from grid cells. It was long theorized [55, 30] and recently been experimentally established [43] that the population activity of grid cells lives on a toroidal manifold. In this case, the encoding stimulus space has two circular coordinate systems—one each for the meridional and longitudinal axes (Fig 4(a))—which can be detected using persistent homology (Fig 4(b)). We denote the longest bar by *θ*_1_ and the second-longest by *θ*_2_. As in the case of our first set of experiments, we initialized a series of three-layer networks (Figure 1) and conducted standard propagation trials for each *θ*_*i*_ to first asses how effectively random connectivity would propagate this pair of circular coordinate systems. For 256 of 360 (71.11%) of network architectures tested, mean PS over 10 trials for each *θ*_*i*_ was below 0.25.

**Figure 4.**
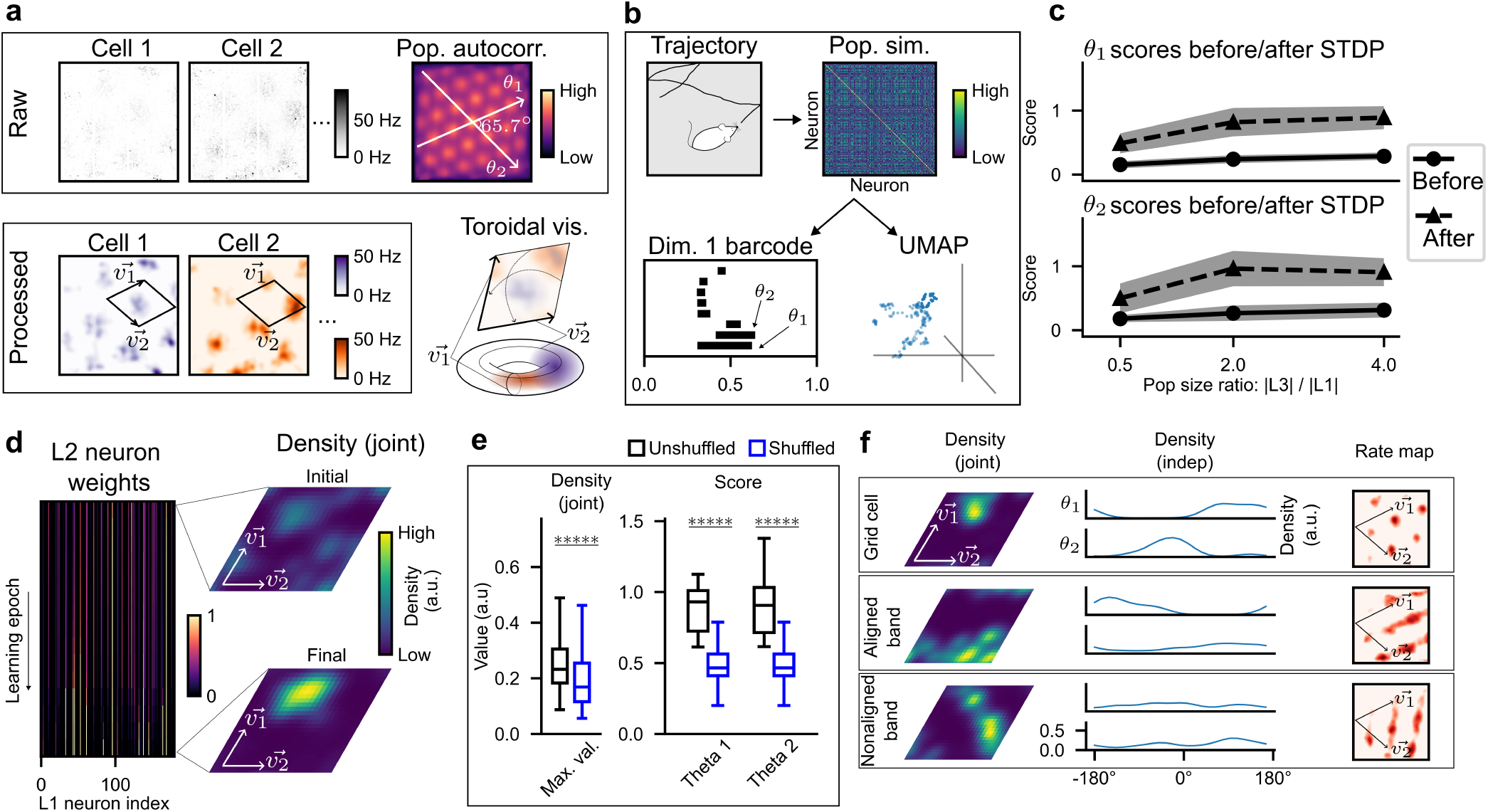
Learning multiple simultaneously encoded circular coordinate systems from grid cells. **(a)**, (Top) Minimally-processed rate maps of grid cells are extracted as in [43] and autocorrelation of each rate map is computed and averaged across the population. Visualization suggests two periodic coordinate systems, *θ*_1_ and *θ*_2_. (Bottom) Rate maps are smoothed and normalized to a maximum firing rate of 50 Hz. Each rate map can be visualized as a single convex receptive field on a toroidal manifold. **(b)**, A simulated trajectory of 25 minutes of freely moving behavior is presented to a collection of 180 synthetic grid cells, and intrapopulation similarity is computed. The barcode reveals two significant circular coordinate systems; UMAP projection fails to identify the toroidal manifold. **(c)**, Hebbian STDP induces propagation of both circular coordinate systems; PS generally increases with L3 population size. **(d)**, (Left) synaptic weights from L1 population to a fixed L2 neuron during learning. (Right) Kernel density estimation of a synaptic-weight-defined random variable on the toroidal manifold is shown before and after learning. A higher maximum value after learning indicates that synaptic weights have concentrated in a localized region of the toroidal manifold. **(e)** A geometric ablation study is performed by shuffling synaptic weights of L2 neurons and average maximum density and PS for each *θ*_*i*_ is recomputed. Shuffling significantly reduces average maximum density and PS for each *θ*_*i*_ (Welch’s t-test *p* < 1 ×10^−5^ for all comparisons). **(f)**, Emergence of several classes of rate maps is observed. Synaptic weight concentrated in (top) a convex region in the toroidal manifold produces a grid cell; (middle) a band parallel to 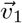 in the toroidal manifold produces a rate map composed spatially periodic bands parallel to 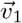; (bottom) a band not aligned to either 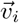 in the toroidal manifold produces a rate map of spatially periodic bands not parallel to either 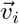.

Interestingly, the effect of inhibitory-bias was not as pronounced as in the case of head-direction cells: 89 of 120 inhibitory-biased architectures showed the same propagation failure but the effect of an *excitatory* bias was profound: in all 120 excitatory-biased architectures, mean PS of each *θ*_*i*_ was below 0.25. These results suggest that random connectivity is often insufficient to propagate multiple circular coordinate systems, and that both inhibitory and excitatory bias exacerbate this failure. For each of the 360 network architectures, we next implemented the same STDP-learning protocol as in the head-direction experiments, using instead simulated free behavior in an open field environment to drive the input layer.

In total, 223 of 360 (61.94 %) architectures showed significantly higher mean PS (paired t-test *p* < 0.05 for each *θ*_*i*_) over 10 trials after the STDP-learning protocol. However, the overall efficacy of STDP-learning was somewhat dampened compared to the head-direction case: only 27 of 360 architectures had mean PS over 0.75 for each *θ*_*i*_ and 103 of 360 had mean PS over 0.5 after STDP-induced learning. The failure of STDP to induce multiple circular coordinate systems downstream was particularly pronounced among the excitatory-biased networks: none had mean PS above 0.75, and only 7 of 120 had mean PS above 0.5 after learning. These results suggests that STDP can induce learning of multiple simultaneously-present circular coordinate systems, but that it is less effective than in the case of a single circular coordinate system and particularly sensitive to changes in excitatory/inhibitory balance. As in the single-coordinate case, we then asked: what other factors determine the efficacy of STDP-learning?

As in our first set of experiments, we fit a decision tree model to predict post-learning PS (averaged across *θ*_1_ and *θ*_2_) based on the seven model hyperparameters and obtained a reasonable fit (*R*^2^ = 0.40). The model revealed three key determining factors: excitatory/inhibitory balance (Δ^2^ = 0.42) as expected from the analyses above, population size ratio |*L*3| */* |*L*1| (Δ*R*^2^ = 0.31), and connection density Δ*R*^2^ = 0.14). Notably, the effect of these hyperparemeters was respectively greater, less than, and greater than the corresponding effect in the case of a single circular coordinate system. Computing Spearman’s rank correlation across all trials for each *θ*_*i*_ showed significant (*p* < 0.05) positive correlation between population size ratio and post-learning PS for 86 of 120 (71.67 %) hyperparameter combinations.

Together, these results suggest that population size plays a key role in the efficacy of STDP-learning, but that excitatory/inhibitory balance is a more significant factor, and that connection density plays a larger role than in the case of a single circular coordinate system.

Given the similarities in overall PS results to the head-direction experiments, we next investigated whether a similar process of weight localization through learning occurred in the grid-cell experiments.

### Local toroidal geometry and the emergence of band cells

To determine the extent to which L2 neurons localized weights on the input toroidal manifold, we performed a similar analysis to the head-direction case by assigning L1 neurons a fixed position on a torus *T* and defining, for each L2 neuron *i* a random variable *X*_*i*_ on *T*. We assessed the localization of synaptic weight by computing a probability density function *f*_*i*_ of *X*_*i*_.

Fig 4(d) shows a representative L2 neuron. Before STDP-learning, synaptic weight is approximately uniformly distributed among L1 neurons on *T*. Through the learning process, weights concentrate in a local neighborhood of *T* (Fig 4 (d) right). We performed a geometric ablation study by shuffling connection weights of L2 neurons and recomputed PS for *θ*_1_ and *θ*_2_. The ⟨max *f*_*i*_⟩ decreased (as designed) and we observed a corresponding decrease in PS for each *θ*_*i*_ (Fig 4(e)). This suggests that the concentration of synaptic weight to local neighborhoods of the toroidal stimulus space plays a key role in propagating circular coordinate systems downstream.

To determine how this reorganization affects tuning properties of downstream neurons, we computed rate maps for L2 and L3 neurons analytically and compared the resultant rate maps to the computed *f*_*i*_ for each neuron. In many cases *f*_*i*_ showed a well-defined peak in a roughly symmetric circular neighborhood of the input torus (Fig 4(f) top row). The corresponding rate map was consistent with a stereotypical grid cell. Remarkably, we also observed a number of other “cell-types.” For instance, if a downstream L2 neuron *i* learned only a *single* circular coordinate system, then *f*_*i*_ formed a single “band” of synaptic weight aligned with one axis of the input torus (Fig 4(f) second row). In this case, the rate map showed activity at spatially periodic bands across the environment. We also observed a number of downstream cells with spatially periodic bands that were not aligned with either *θ*_1_ or *θ*_2_ (Fig 4(f) third row). Cells characterized by these spatially periodic bands were first reported [44] and explored further in [56]. Our results suggests that these cells may emerge naturally as a downstream effect of spike-timing dependent plasticity in the context of grid-like input.

## Discussion

In this study, we identified a simple biophysical mechanism — Hebbian spike-timing dependent plasticity — that can induce learning of circular coordinate systems in feedforward networks. We determined that random connectivity fails to propagate circular or toroidal topology in a broad range of connectivity parameters, and identified key factors that govern the efficacy of Hebbian STDP to induce topological learning. A common theme emerged across conditions: downstream neurons reorganize synaptic weight to localized neighborhoods of the input stimulus space. Notably, this mechanism of learning local geometric structure led to the spontaneous formation of bimodally-tuned head direction cells and periodic band-like receptive fields in downstream neurons. These results highlight how simple biophysical mechanisms can induce learning of nonlinear neural manifolds while simultaneously producing complex neural codes and new emergent geometric features similar to those found in real neural systems.

Many studies have used tools from topological data analysis to uncover underlying stimulus space structure [57, 58, 59, 60, 61, 43, 62]. There is also a wide literature on learning manifold structure using techniques from machine learning, including gradient back-propagation. In this study, we asked a different question: what happens when one combines complex stimulus space topology with a simple biophysical mechanism to update synaptic weights? Our application of recently developed methods from topological data analysis was vital to our ability to quantitatively evalute the efficacy of this learning process.

Our focus on circular coordinate systems was motivated by their prevalence in observed neural systems, and facilitated by the tractability of identifying and tracking the topology of the associated stimulus space using tools of topological data analysis. However, the identified mechanism by which Hebbian STDP reorganizes feedforward architectures in the presence of input topology suggests a general phenomenon. In this sense, there is nothing special about circular or toroidal stimulus spaces: we expect that the same mechanism may be used to induce stimulus space structure in downstream populations with arbitrary topology. However, detecting and quantitatively matching coding schemes associated to arbitrary neural manifolds would require extensions to our current analytic techniques and is left for future investigation.

In summary, we have shown that Hebbian STDP is sufficient to induce learning of a single or multiple simultaneously encoded circular coordinate systems. Our results reveal a common mechanism underlying the learning of local geometry and global topology that suggests a universal principle by which stimulus space structure may be learned. As the information encoded by many neural systems, including head-direction cells, simple cells, and grid cells, are topologically structured, we posit that a better understanding of how biophysical mechanisms interact with such structure will provide novel insights into the functioning of these systems.

## Materials and Methods

### Experimental data

Raw data for the head-direction experiments consisted of extracellular recordings from multi-site silicon probes in the anterior thalamus and subicular formation of freely moving mice and was acquired from the published data sets in [2, 42]. Cells during awake exploration of Mouse 12, Mouse 17, Mouse 20, Mouse 24, Mouse 25, and Mouse 28 were used. Raw data for the grid cell experiments were acquired from the published data of [43] and associated Jupyter notebooks. These data consisted of Neuropixel recordings targeting the MEC–parasubiculum (PaS) region in three Long Evans rats. We used only the data of Rat R during open-field exploration.

### Extraction and processing of tuning curves and rate maps

For the first set of experiments (head-direction) tuning curves for 969 neurons were computed by averaging firing rate as a function of head direction. Three synthetic tuning curves were generated from each empirical curve by rotating by 90°, 180°, and 270°, yielding a population of 3876 synthetic L1 neurons. To ensure all neurons (those well-tuned to the circular stimulus, and those not well-tuned) contributed on roughly equal footing to the L1 population activity, tuning curves were normalized to a uniform maximum firing rate of 50 Hz. This rate was near the upper bound of observed peak firing for neurons well-tuned to head direction, while neurons not well-tuned generally fired at lower rates.

For the second set of experiments (grid cells) rate maps of 166 neurons were computed by averaging firing rate as a function of location during ~122 minutes of free behavior in a square enclosure using the published software associated to [43]. Computing the average autocorrelation of the neural population revealed prominent grid structure and two basis vectors 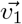 and 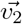 for the grid were chosen by aligning axes to local maxima of the average autocorrelation (Fig 4(a)). Next, 36 synthetic rate maps were generated from each empirical rate map by shifting along the vector 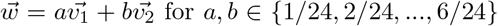. This yielded a population of 5976 synthetic L1 neurons. Rate maps were then mean-subtracted and thresholded and normalized to a maximum firing rate of 50 Hz.

### Network initialization and architecture

We modeled a feed-forward neural network with three layers, L1, L2, and L3 (see Fig 1(b)). L1 neurons were modeled as excitatory Dale neurons with positive outgoing weights and the L2 to L3 architecture was fixed so that each L3 neuron received inputs from a unique pair of L2 neurons—one with weight +1 (excitatory) and one with weight −1 (inhibitory). Consequently, each network satisfied |L2| = 2 |L3|. For the head-direction experiments, |L1| was chosen from 60, 90, 120 and for the grid cell experiments, L1 was chosen from {120, 180}.

Excitatory/inhibitory bias was introduced by drawing L1 to L2 weights from normal distributions *D* with negative, zero, or positive means, specifically 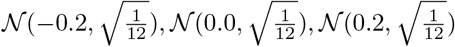. Standard deviations were chosen to match that of the standard uniform distribution *U* [0, 1]. For each fixed L1 neuron, a weight *w* ~ *D* was drawn and a synapse of magnitude *w* was created between *i* and a randomly chosen L2 neuron: excitatory if *w* ≥0, inhibitory if *w* < 0. This process was repeated until the specified connection density was reached. For the head-direction experiments, connection densities for the |L|1 to |L2| layer were chosen from{0.25, 0.5, 0.75} and for the grid cell experiments from {0.15, 0.20, 0.25, 0.50, 0.75}.

Noise in the L1 encoding of circular variables was introduced as follows. For the head-direction experiments, we computed the Skaggs information index [63] of each tuning curve, which is high for neurons with unimodal tuning curves and low for tuning curves not well-correlated to heading. Tuning curves for L1 neurons were then sampled according to one of several empirical distributions of Skaggs index. For example, given the distribution [0, 0.1, 0.2, 0.3, 0.4], 40% of L1 neurons were chosen with Skaggs index in the top 20% of the population of 3876 neurons, 30% were chosen with Skaggs index in the 60^th^ − 80^th^ percentile, and so on. For the head-direction experiments, we sampled from three distributions ranging from no noise to high levels of noise:

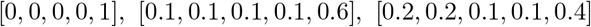

The grid cell experiments were handled similarly but using the “gridness” score of [64] in place of Skaggs index. For these experiments, we sampled from two distributions of gridness scores: [0, 0, 0, 0, 10] and [0, 0, 1, 1, 8].

### Calculating downstream response

In all experiments, each projection L1→L2 and L2→L3 was modeled as a one-layer threshold-linear feedforward rate network. The instantaneous firing rate *r*(*i*) of a downstream neuron *i* was given by

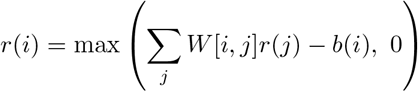

where *r*(*j*) is the instantaneous firing rate of upstream neuron *j, W* is the synaptic weight matrix and *b*(*i*) is the bias term. Rate-dependent Gaussian noise *N* was added to each *r*(*i*), with *N* = max (0, 𝒩 (*r*(*i*), *σ*^2^)) and *σ* = max(0.05*r*(*i*), 2.5).

Each L2 neuron *i* was assigned an adaptive bias *b*(*i*) set as half the maximum firing rate neuron *i* had exhibited during the simulation; this is consistent with observed mechanisms for homeostatic control of intrinsic excitability [65, 66, 67, 68]. Thresholds were initialized either uniformly at 0 or at a “pre-trained” value determined by simulating 10s of stimulus and setting *b*(*i*) to half the maximum firing rate observed for neuron *i* over this warm-start epoch. Layer L3 was reserved as a readout layer and, hence, its neurons were assigned fixed thresholds *b*(*i*) = 0.

### Heading and free behavior simulations

To simulate realistic heading stimuli for the head-direction experiments, the tracked headings of 6 mice over a total of 31 trials of open-field exploration sessions were analyzed. The changes in heading angle (radians) per 10ms bin over the entire set were well-approximated by a lognormal distribution with parameters *µ* =−4.390 and *σ* = 1.181. The probability that the heading continued in the same direction (i.e., clockwise or counterclockwise) between consecutive 10ms bins was calculated to be *p* = 0.852. The simulated heading stimulus was then modeled as a biased random walk on a circle, each point on the walk corresponding to an angular coordinate in [0, 2*π*), with step size and directional bias chosen to match the data above. Finally, the resulting simulated trajectories were visually confirmed to approximate the experimental data well.

The simulated free-behavior for the grid cell experiments was modeled similarly. In this case, the changes in heading angle per 10ms bin of the empirical heading time series were well-approximated by a lognormal distribution with parameters *µ*_1_ =− 5.091 and *σ*_1_ = 1.434 capped at a maximum of 0.164, equal to that of the experimental data.

Next, trajectory displacements greater than a threshold *τ* = 0.6mm over 10ms bins were collected and approximated by a normal distribution with mean *µ*_2_ = 0.00165 and standard deviation *σ*_2_ = 0.000817. Samples were drawn from the fitted distribution, with a minimum of zero enforced for displacement and visually confirmed via histogram comparison to match the experimental data. The threshold *τ* was selected heuristically to ensure visual consistency with the experimental distribution. Finally, free-behavior was modeled as a random walk in a square enclosure with step size and heading drawn from the fitted distributions. On collision with a boundary of the environment, a new heading was uniformly sampled from [0, 2*π*).

### STDP updates and normalizations

The same STDP and normalization protocol was used for both the head-direction and grid cell experiments. Over a 1-second learning epoch (discretized in 10ms bins), instantaneous firing rates of L1 neurons were computed according to their tuning curves (resp. rate maps) and Gaussian noise proportional to the firing rate was added. Rates of L2 neurons were computed as a rectified linear combination of input signals with a linear threshold term. Firing rates were converted to spike trains by sampling an inhomogeneous Poisson process. To model synaptic transmission delay, spiketrains of L2 neurons were uniformly shifted by a randomly chosen delay between 1ms and 9ms (inclusive).

Synaptic weights were then updated by applying the spike-timing dependent plasticity (STDP) equations in [39, Fig 1] to the respective presynaptic and postsyaptic spiketrains using a±50ms search window centered on each presynaptic spike (Fig 2(b)). In the event of multiple postsynaptic spikes occurring in the search window, only that nearest in time to the presynaptic spike was considered. To amplify plasticity of synapses involving presynaptic neurons with higher variability in firing rates, these weight changes were then scaled as follows. For a given L2 neuron, STDP-updates result in a set of synaptic updates {Δ*w*_*i*_}_*i* ∈*I*_. For each *i*, let 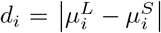 where 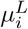 and 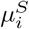 are the mean firing rates of presynaptic neuron *i* over the previous 10s and during the considered 1s epoch, respectively. Let *m* = min_*i*_ {*d*_*i*_}, *M* = max_*i*_ {*d*_*i*_}, and 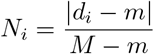. Each Δ*w*_*i*_ was then scaled by *N*_*i*_. Additionally, weights projecting to each L2 neuron were normalized to enforce a maximum weight of 1 and to preserve total incoming synaptic weight. This update was repeated over 5,000 learning epochs for a total learning time of 5,000s or approximately 83 minutes.

### Propagation Scores

The Propagation Score (PS) of a fixed network was calculated by simulating 25 minutes of stimulus input—heading trajectory for head-direction experiments and free behavior for grid cells. Responses of L1 and downstream neurons were computed as described above. To introduce additional variability and reduce dependence on precise rate profiles, firing rates of L1 and L3 neurons were converted to spiketrains by sampling an inhomogeneous Poisson process and then back to rates by binning and smoothing. Next, dissimilarity matrices *D*_1_ and *D*_3_ of size |L1| *×* |L1| and |L3| *×* |L3|, respectively, were computed using cross-correlograms (as in [59]) of firing rate rasters for the L1 (resp. L3) populations. An |L1| *×* |L3| cross-population dissimilarity matrix *D*_13_ was similarly computed.

Filtered clique complexes were constructed from *D*_1_ and *D*_3_, and a filtered witness complex was constructed from *D*_13_. The associated barcodes *BC*_1_, *BC*_3_, and *BC*_13_ were computed and, to assess downstream learning of the topological feature(s) encoded by the L1 population, the method of analogous cycles [32] was used to detect matches between features in *BC*_1_ and *BC*_3_. As detailed in [32], a match between bars implies that they represent the same information—in our case, the same circular coordinate system. In the context of filtered clique complexes, the lifetime (or persistence) of a bar is correlated with its robustness as a feature of the data. This means that long-lived (high-persistence) bars correspond to robust circular coordinates systems in the encoding. For the head-direction experiments, the single highest-persistence bar in *BC*_1_ was compared to all features in *BC*_3_; for grid-cells, each of the two highest-persistence bars in *BC*_1_ was compared similarly. If matches were found, PS was calculated as the persistence of the match in *BC*_3_ divided by the persistence of the selected feature in *BC*_1_.

### Density estimations

To compute the density function of an L2 neuron *i* (Fig 2(a), (e)) for the head-direction experiments, each L1 neuron was assigned a position on the circle *S*^1^ = [0, 2*π*) by the location of the maximum value of its computed tuning curve. If this location was not unique, we chose the location corresponding to smallest angle between 0° and 360° by convention. An empirical distribution on *S*^1^ = [0, 2*π*) was constructed by sampling the circle-valued positions of L1 neurons that projected to neuron *i*, with each position replicated at a rate proportional to synaptic weight. The probability density function of this distribution was then approximated using kernel density estimation with a von Mises kernel (parameter *κ* = 20 and 100 bins). This approximation was used as the density function of neuron *i*. Density functions of L2 neurons in the grid cell experiments were similarly computed: L1 neurons were assigned a position on *T* = *S*^1^ ×*S*^1^ by the location of their rate map maximums, and the probability density function of an analogously defined distribution on *T* was approximated using a product of von Mises kernels—one for each circular coordinate in the toroidal encoding.

## Acknowledgements

The authors thank Ben Dunn and Erik Hermansen for sharing code to extract tuning curves and rate maps from the raw data used in all experiments; Carina Curto, Kathryn Hess, Nicole Sanderson, and Iris Yoon for helpful feedback throughout the project; and Gregory Henselman-Petrusek for assistance developing software using the Open Applied Topology package. Both authors were partially supported by the Air Force Office of Scientific Research under award number FA9550-21-1-0266. Both authors began this research while appointed at the University of Delaware.

## Notes

### Competing Interest Statement

The authors have declared no competing interest.

